# Genomic risk prediction of coronary artery disease in nearly 500,000 adults: implications for early screening and primary prevention

**DOI:** 10.1101/250712

**Authors:** Michael Inouye, Gad Abraham, Christopher P. Nelson, Angela M. Wood, Michael J. Sweeting, Frank Dudbridge, Florence Y. Lai, Stephen Kaptoge, Marta Brozynska, Tingting Wang, Shu Ye, Thomas R Webb, Martin K. Rutter, Ioanna Tzoulaki, Riyaz S. Patel, Ruth J.F. Loos, Bernard Keavney, Harry Hemingway, John Thompson, Hugh Watkins, Panos Deloukas, Emanuele Di Angelantonio, Adam S. Butterworth, John Danesh, Nilesh J. Samani, for The UK Biobank CardioMetabolic Consortium CHD Working Group

## Abstract

**Background:** Coronary artery disease (CAD) has substantial heritability and a polygenic architecture; however, genomic risk scores have not yet leveraged the totality of genetic information available nor been externally tested at population-scale to show potential utility in primary prevention.

**Methods:** Using a meta-analytic approach to combine large-scale genome-wide and targeted genetic association data, we developed a new genomic risk score for CAD (metaGRS), consisting of 1.7 million genetic variants. We externally tested metaGRS, individually and in combination with available conventional risk factors, in 22,242 CAD cases and 460,387 non-cases from UK Biobank.

**Findings:** In UK Biobank, a standard deviation increase in metaGRS had a hazard ratio (HR) of 1.71 (95% CI 1.68–1.73) for CAD, greater than any other externally tested genetic risk score. Individuals in the top 20% of the metaGRS distribution had a HR of 4.17 (95% CI 3.97–4.38) compared with those in the bottom 20%. The metaGRS had higher C-index (C=0.623, 95% CI 0.615–0.631) for incident CAD than any of four conventional factors (smoking, diabetes, hypertension, and body mass index), and addition of the metaGRS to a model of conventional risk factors increased C-index by 3.7%. In individuals on lipid-lowering or anti-hypertensive medications at recruitment, metaGRS hazard for incident CAD was significantly but only partially attenuated with HR of 2.83 (95% CI 2.61– 3.07) between the top and bottom 20% of the metaGRS distribution.

**Interpretation:** Recent genetic association studies have yielded enough information to meaningfully stratify individuals using the metaGRS for CAD risk in both early and later life, thus enabling targeted primary intervention in combination with conventional risk factors. The metaGRS effect was partially attenuated by lipid and blood pressure-lowering medication, however other prevention strategies will be required to fully benefit from earlier genomic risk stratification.

**Funding:** National Health and Medical Research Council of Australia, British Heart Foundation, Australian Heart Foundation.

## Introduction

Coronary artery disease (CAD) is the leading cause of morbidity and mortality worldwide, and early identification of individuals at high risk of CAD is essential for primary prevention. While conventional CAD risk factors, such as lipids, blood pressure, and smoking, become predictive in middle life, their predictive ability is weaker at a younger age. The heritability of CAD has been estimated to be 40– 60% and thus genetic predisposition is a risk factor of significant potential for earlier risk prediction ^1,2^. Over the last 10 years, genome-wide association studies have begun elucidating the genetic architecture of CAD and laid the foundation for developing genomic risk scores (GRSs) for estimating an individual’s underlying genomic risk ^3-9^. However, previous GRSs for CAD have not facilitated fundamental change in early CAD risk screening strategies as they have not reached a sufficient level of predictive power, e.g. by outperforming conventional cardiovascular risk factors, nor have they established general applicability via large-scale external testing in representative population-based samples. A likely reason is that the polygenicity of CAD has not been sufficiently reflected in the predictive genetic models, together with imprecision in the effect size estimates driven by limited sample sizes ^10-12^. Previously published GRSs have utilised only genetic variants of genome-wide significance ^4,5,8^ or were based on arrays that focus on pre-selected loci ^3^; thus, they have not fully utilised genome-wide variation, and have not been able to accurately estimate the relative contribution of each genetic variant to CAD risk. Furthermore, the generalisability of previous GRSs has been limited by lack of external testing in truly large-scale cohorts that represent a diversity of ancestries ^3,13^, and a wide spectrum of the CAD burden, e.g. not only myocardial infarction ^14,15^. A more powerful and generalisable genome-wide GRS for CAD would likely have far reaching implications for early screening at a population level, prioritisation for lifestyle and therapeutic intervention, and targeted clinical trials.

Here, we utilise a meta-analytic strategy to construct a GRS for CAD (metaGRS) that captures the totality of information from the largest previous genome-wide association studies, and then investigate the external performance of this metaGRS in stratifying CAD risk in >480,000 individuals from the UK Biobank (UKB) ^16^. Furthermore, we assess the effects of several conventional risk factors (smoking, blood pressure, BMI, diabetes) on different genomic risk backgrounds, with the aim of identifying subsets of individuals who are likely to benefit from earlier and more intensive screening, or who may not benefit from screening until later life. Finally, to assess the potential therapeutic implications of genomic risk scores, we test the impact of blood pressure and lipid lowering medication on the performance of metaGRS.

## Methods

### Study design and participants

Details of the design of the UKB have been reported previously ^12^. Participants were members of the UK general population aged between 40–69 years at recruitment, identified through primary care lists, who accepted an invitation to attend one of the 22 assessment centers that were serially established across the UK between 2006 and 2010. At recruitment, detailed information was collected via a standardised questionnaire on socio-demographic characteristics, health status and physician-diagnosed medical conditions, family history and lifestyle factors. Selected physical and functional measurements were obtained including height, weight, waist-hip ratio, and systolic and diastolic blood pressures. The UKB data were subsequently linked to Hospital Episode Statistics (HES) data, as well as national death and cancer registries. The HES data available for the current analysis covers all hospital admissions to NHS hospitals in England and Scotland from April 1997 to March 2015, with the Scottish data dating back as early as 1981. HES uses International Classification of Diseases ICD 9 and 10 to record diagnosis information, and OPCS-4 (Office of Population, Censuses and Surveys: Classification of Interventions and Procedures, version 4) to code operative procedures. Death registries include all deaths in the UK up to January 2016, with both primary and contributory causes of death coded in ICD-10.

CAD was defined as fatal or non-fatal myocardial infarction (MI) cases, percutaneous transluminal coronary angioplasty (PTCA), or coronary artery bypass graft (CABG). The age of event in prevalent cases was determined by self-reported age and calculated age based on the earliest hospital record for the event; if both self-reported age and calculated age were available, the smaller value was used. For incident cases, hospital and/or death records were used to determined age of event. Prevalent vs incident status was relative to the first UKB assessment. In UKB self-reported data, cases were defined as having heart attack diagnosed by doctor (data field #6150) or ‘non-cancer illnesses that self-reported as heart attack’ (data field #20002) or self-reported operation including PTCA, CABG, or triple heart bypass (data field #20004). In HES hospital episodes data and death registry data, MI was defined as hospital admission or cause of death due to ICD9 410–412, ICD10 I21–I24, or I25.2; CABG, PTCA were defined as hospital admission OPCS-4 K40–K46, K49, K50.1, or K75.

We defined risk factors at the first assessment as follows: diabetes diagnosed by doctor (field #2443), body mass index (BMI; field #21001), current smoking (field #20116), and hypertension. For hypertension we used an expanded definition including self-reported high blood pressure (either on blood pressure medication, data fields #6177, #6153; or systolic blood pressure >140 mmHg, fields #4080, #93; or diastolic blood pressure >90 mmHg, data fields #4079, #94). For the analyses of the number of elevated risk factors, we considered diagnosed diabetes (Y/N), hypertension at assessment (Y/N), BMI >30 kg/m^2^, and smoking at assessment (Y/N).

Genotyping of UK Biobank participants was undertaken using a custom-built genome-wide array (the UK Biobank Axiom array: http://www.ukbiobank.ac.uk/wp-content/uploads/2014/04/UK-Biobank-Axiom-Array-Datasheet-2014.pdf) of ~826,000 markers. Genotyping was done in two phases. 50,000 subjects were initially typed as part of the UK BiLEVE project ^13^. The rest of the participants were genotyped using a slightly modified array. Imputation to ~92 million markers was subsequently carried out using the Haplotype Reference Consortium (HRC) ^17^ and UK10K/1000Genomes haplotype resource panels, however at the time of analysis, known issues existed with the imputation using the latter panel.

### Data processing and quality control

Only autosomal genetic variants imputed using the HRC panel and which had MAF >0.1% were included in our analyses, totalling 14.5 million variants. We converted the imputed dosages to PLINK^18^ genotype calls with minimum probability 0.9 (otherwise the call was set to missing; we removed variants with >1% missingness across individuals). To control for population structure, we utilised the genetic principal components (PCs) as given by the UKB^19^.

Two QC schemes were used. For the GRS46K and FDR202 genetic risk scores (defined below), we kept variants with MAF >0.1%, however filtering by HWE and imputation quality (INFO) was not employed as this led to fewer variants mapping to these scores and thus reductions in predictive power. For the 1000Genomes CAD score, we included variants with impute2 INFO >0.01 and HWE *P*>10^−12^. A lenient HWE threshold was used because HWE was computed over all individuals without regard to population structure, thus variants that deviation from HWE at a stringent threshold may be needlessly excluded as they may not be indicative of genotyping error. A lenient INFO threshold (e.g. relative to the INFO >0.4 used in Nikpay et al ^20^) was employed as the UKB has a relatively large sample size, and an analysis of variants with INFO=0.001 in the UKB has equivalent statistical power to an analysis of 0.001 × 500,000 = 500 individuals; thus genetic variants with low INFO score can still offer improvements in predictive accuracy. Similarly, bias in odds ratios arising from genotyping or imputation error has the effect only of reducing the predictive accuracy of a GRS, but this reduction is less than would occur by omitting the marker entirely. After the above quality control procedures were implemented, 14,516,436 autosomal markers were available for subsequent analyses. For the 1000Genomes CAD score, we only used markers found in both the UKB data and the 1000Genomes CAD summary statistics, resulting in 5,176,852 autosomal variants. In the UKB, 485,629 individuals had matching genetic data and CAD outcome data. We removed individuals with (i) diagnoses in one of ICD9 414.1, ICD10 I25.0, I25.3, I25.4 or (ii) CAD event but no known age or date of CAD. There was no evidence of differential missingness in genetic data between CAD and non-CAD individuals. The final dataset consisted of n=485,629 individuals, including 22,242 CAD events and 460,387 non-cases.

### Construction of metaGRS

To construct a meta genomic risk score (metaGRS) for CAD, we followed a meta-analysis strategy using the largest CARDIoGRAMplusC4D genome-wide association analysis without UKB data ^20^ and the CARDIoGRAMplusC4D Metabochip analysis, which focused on variants and regions known or thought to be associated with cardiometabolic phenotypes ^21^. The Metabochip-based GRS46K (46,000 genetic variants, excluding ~3,000 A/T and C/G SNPs) was constructed previously ^3^, while the FDR202 and ‘1000Genomes’ CAD GRS derived from the CARDIoGRAMplusC4D genome-wide association analysis, are described below. The three GRSs all provide an imperfect measure of an individual’s genomic risk of developing CAD, due to incomplete and targeted coverage of the genome, finite sample estimates of the marginal effect sizes for each genetic variant, and genotyping or imputation uncertainty. Since it is well known that a risk factor measured with error can attenuate the association between the risk factor and disease occurrence (regression dilution bias^22^), we reasoned that a ‘meta score’, the weighted average of the three standardised genetic risk scores, would provide a more precise estimate of an individual’s genomic risk of developing CAD.

To create the metaGRS, we used the meta-analysis summary statistics ^20^, consisting of dbSNP rsid, risk allele, and effect size (log odds ratio). We used the existing GRS46K ^3^ and FDR202 ^20^ scores. For the GRS46K, we could map 45,810 variants (98%) in the UKB imputed dataset; for the FDR202, 198 variants or proxies thereof mapped (98%; for proxies, minimum *r*^2^ in 1000Genomes CEU+GBR was 0.68, the median *r*^2^ was 1.0).

Each score *s* is the sum of the minor allele dosages of each variant multiplied by its marginal effect size *β* (log odds ratio per dosage of minor allele):

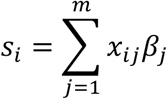

where *x_ij_* ∈ {0, 1, 2} is the count of the minor allele for the jth variant in the ith individual.

To derive a new genomic risk score based on the CARDIoGRAMplusC4D 1000Genomes-imputed GWAS summary statistics ^20^, we split the UKB data into a training set (n=3000) and a validation set (n=482,629). For the training set, we randomly selected 1000 prevalent CAD cases and 2000 non-CAD individuals. We then used PLINK random linkage disequilibrium (LD) pruning to create different scores, based on SNP sets with varying levels of LD and corresponding CARDIoGRAMplusC4D summary statistics, and evaluated their performance on the n=3000 training set, in terms of hazard ratio (HR) per standard deviation (s.d.) of the score (age-as-time-scale Cox regression, stratified by sex and adjusting for BiLEVE genotyping array and 10 genetic PCs). The score with the highest HR corresponded to the *r*^2^ thinning threshold of 0.9 and consisted of ~1.7 million variants (**Supplementary Figure 1**).

The correlation between the three GRSs was moderate (Pearson’s correlation *r*=0.11 for FDR202-GRS46K, *r*=0.19 for FDR202-1000Genomes, *r*=0.27 for GRS46K-1000Genomes), indicating partial but imperfect overlap of the genetic signals captured by each, likely due to shared genetic loci, LD, and partial overlap of individuals in the cohorts used for deriving these summary statistics^20,21^. Such correlation is accounted for in the weighting below.

We derived a meta score (‘metaGRS’), consisting of a weighted average of the standardised scores

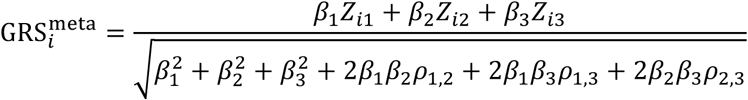

where *Z*_*i*1_, *Z*_*i*2_, *Z*_*i*3_ are the (zero-mean and unit-variance standardised) GRS46K, FDR202, and 1000Genomes CAD risk scores for the *i*th individual, respectively, *β*_1_, *β*_2_, *β*_3_ are the univariate log HRs for each score (estimated using Cox regression in the training set), and *ρ_i,j_* is the Pearson correlation between the *i*th and *j*th scores (in the training set). The univariate log HRs were 0.1278, 0.2359 and 0.2400 per 1-s.d. for the GRS46K, FDR202, and 1000Genomes CAD scores, respectively. To convert this meta-score to a SNP-level score, we used the weighted sum over all *m* = 1,745,180 SNPs (the union of the SNPs in the three scores, and ignoring constant terms),

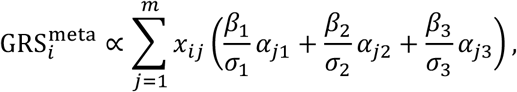

where *σ*_1_, *σ*_2_, *σ*_3_ are the empirical s.d.’s of the scores (GRS46K, FDR202, and 1000Genomes CAD) in the training data, *α*_*j*1_, *α*_*j*2_, *α*_*j*3_ are the SNP effect sizes (log odds ratios from the published summary statistics) for the *j*th SNP in each of the three scores, respectively, and *x_ij_* is the genotype for the *i*th individual’s *j*th SNP. A SNP’s effect size *α_jk_* was considered to be zero for the *k*th score if the SNP was not included in that score.

### Statistical analysis

All scores were standardised to zero-mean and unit-variance. All scores were evaluated using logistic regression or age-as-time-scale Cox proportional hazards regression, with censoring at 75y, as well as with Kaplan-Meier estimates of cumulative incidence (censored at 75y). Unless otherwise noted, analyses using only genetic risk scores include both prevalent and incident CAD cases (germline DNA variation being determined prior to any disease); to avoid reverse causation, analyses that included conventional risk factors (measured at the UKB assessment) used only incident CAD. The Cox models were stratified by sex and adjusted for genotyping array (BiLEVE vs UKB) and 10 genetic PCs. C-indices for the Cox models were sex stratified, using age as time scale. A competing risk analysis, using the Aalen-Johansen estimator (three states: CAD, non-CAD death, and censored), was conducted using the R package ‘survival’^23^.

## Results

The characteristics of the UKB subjects in the external validation set (n=482,629) are shown in Table 1, comprising 22,242 CAD cases before age 75y and 460,387 non-cases in total. There were 9,729 prevalent cases of CAD at the time of recruitment and a further 12,513 incident cases of CAD during a mean follow-up of 6.2 years, at the censoring age of 75 years in 2017. Our meta-analysis approach resulted in a ‘metaGRS’ comprising 1,745,180 genetic variants. A comparison of the metaGRS with its individual components and previously published GRSs from Tikkanen et al ^6^ and Tada et al ^8^ is given in Figure 1, showing the metaGRS had substantially greater association with CAD risk, in terms of hazard ratio as well as positive predictive value (PPV) at any given sensitivity.

**Table 1:**
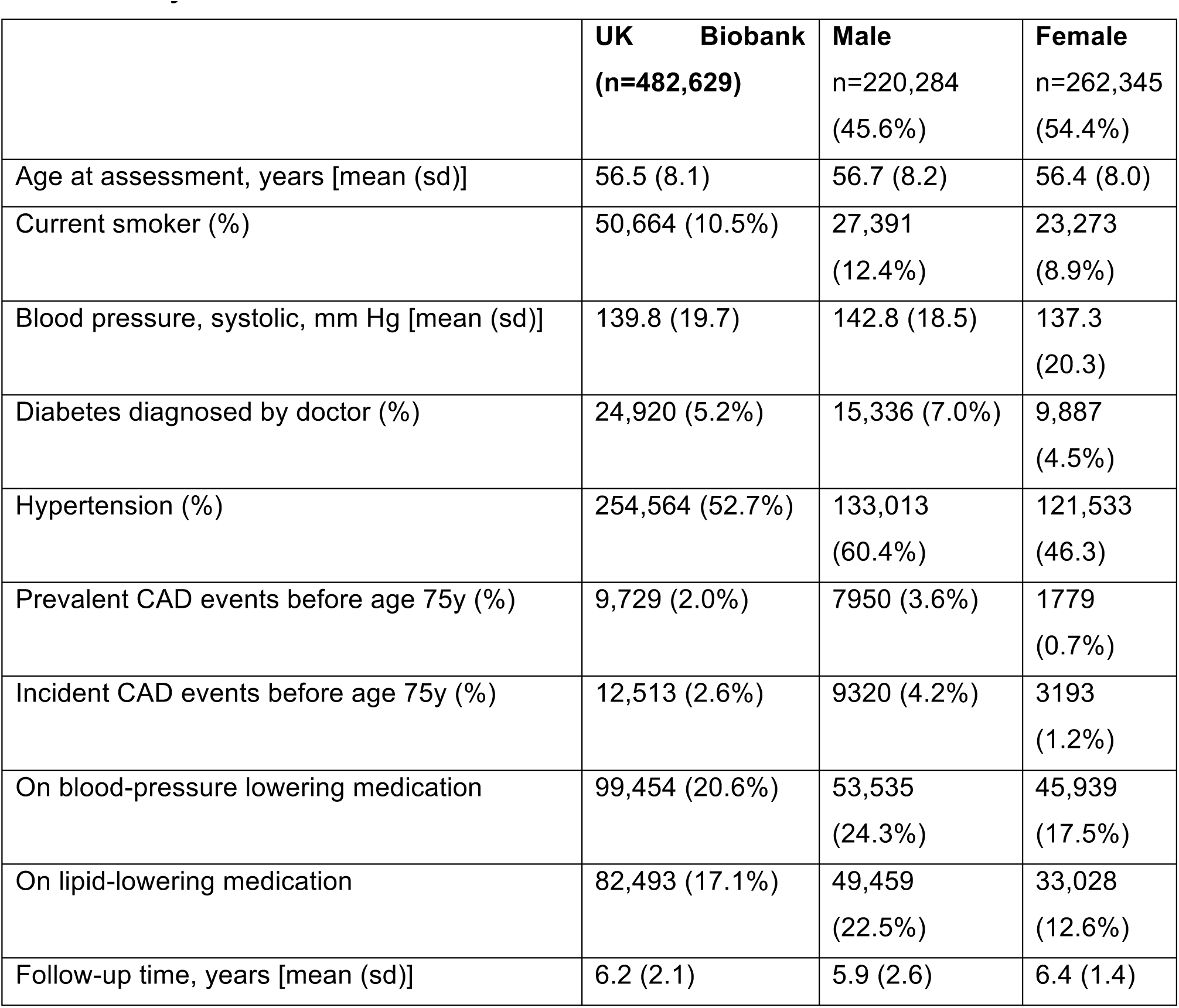
Study characteristics

**Figure 1:**
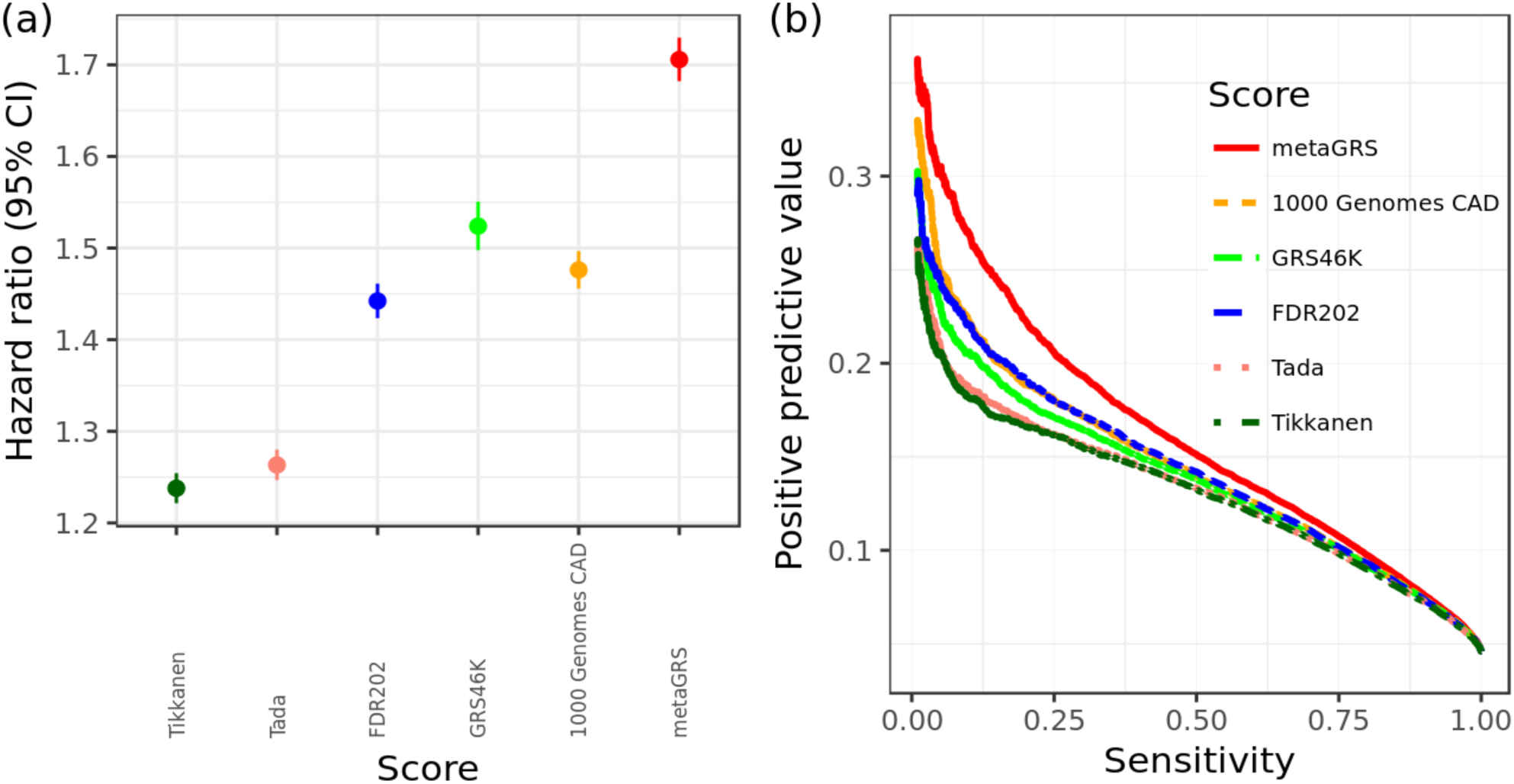
Relative performance of individual genomic risk scores for CAD compared to the metaGRS. In the UKB testing set (n=482,629), (a) hazard ratios per s.d. of each score for all CAD (n=22,242), censored at 75y, from Cox regression stratified by sex and adjusted for genotyping array (BiLEVE/UKB) and 10 genetic PCs; (b) Positive predictive value vs sensitivity for a logistic regression for each GRS, adjusted for sex, age, genotyping array (BiLEVE/UKB) and 10 genetic PCs.

In the external UKB validation set, the metaGRS was accurate at classifying CAD cases vs non-cases with an area under the ROC curve (AUC) of 0.79 (+2.8% over the reference logistic model consisting of sex, age at assessment, genotyping array, and 10 PCs). The metaGRS offered greater PPV at any given sensitivity and thus greater Area under the Precision-Recall Curve (APRC) compared to the reference model (0.161 vs 0.123; Figure 2a). The distributions of the metaGRS amongst prevalent CAD cases, incident CAD cases and non-CAD were each approximately Gaussian and revealed a trend of increasing genomic risk (**Supplementary Figure 2**), with prevalent cases more easily differentiable as they comprise individuals at higher genomic risk who have earlier CAD events.

**Figure 2:**
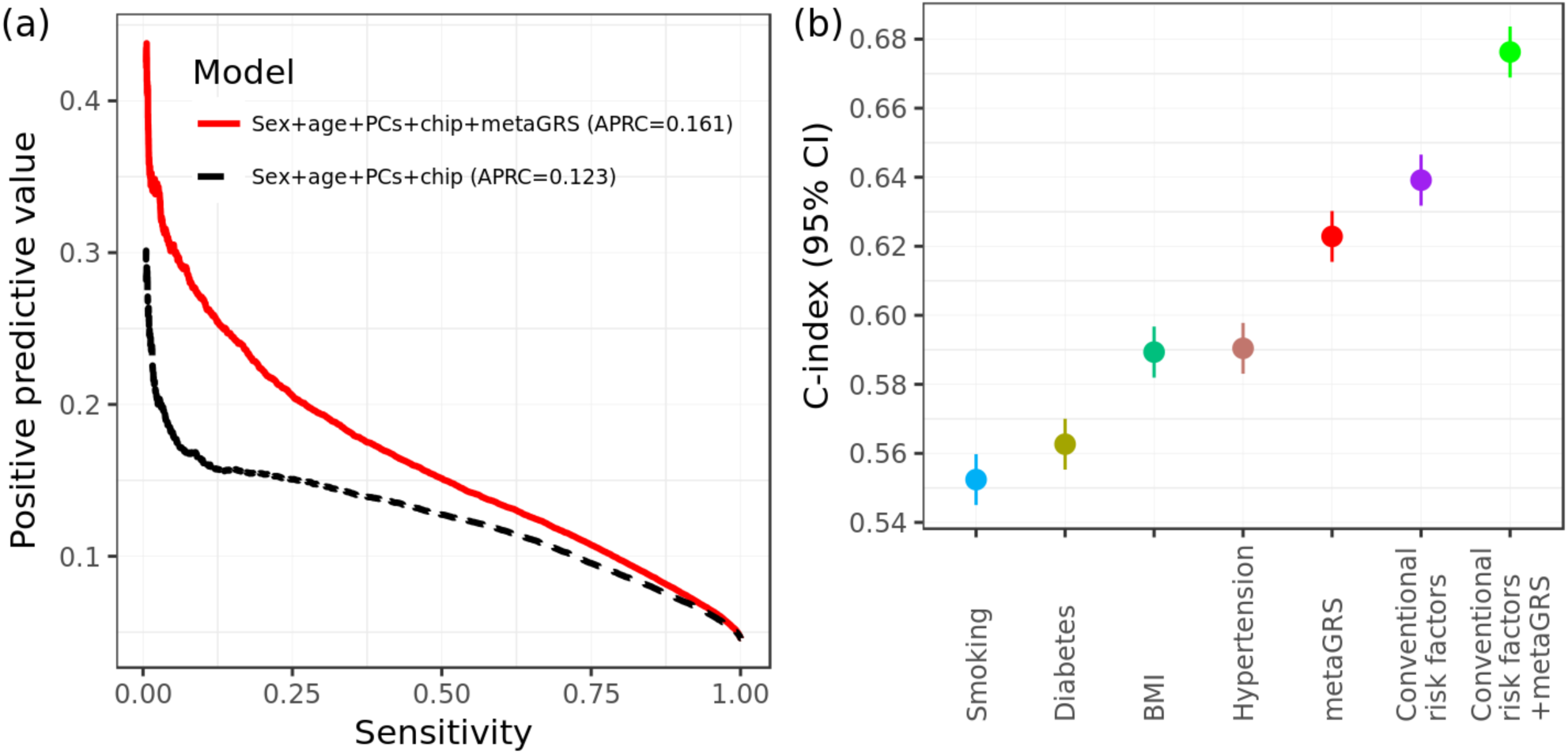
Predictive measures of CAD using the metaGRS and conventional risk factors. (a) Positive predictive values vs sensitivity for the reference model (sex + age + array + 10 genetic PCs) and when adding the metaGRS to the model for all CAD in the UKB testing set. APRC is Area under the Precision-Recall Curve. (b) C-index for sex-stratified age-as-time-scale Cox regression of incident CAD for conventional risk factors individually and in combination with the metaGRS, including genotyping array and 10 genetic PCs as covariates.

In sex-stratified Cox regression models for CAD, the metaGRS had an HR of 1.71 (95% CI 1.68– 1.73) per s.d. of metaGRS (P<0.0001) (Figure 1). The metaGRS was significantly but weakly associated with body mass index (BMI) at assessment (0.0044 log(kg/m^2^) per s.d., 95% CI 0.0039– 0.0049, P<0.0001), diagnosed diabetes (OR=1.14 per s.d., 95% CI 1.13–1.16, P<0.0001), hypertension at assessment (OR=1.19 per s.d., 95% CI 1.18–1.20, P<0.0001), and current smoking at assessment (OR=1.06 per s.d., 95% CI 1.04–1.07, P<0.0001). No evidence for competing risk effects was observed (**Supplementary Figure 3**). In Cox regression of incident CAD (Figure 2b), models based on the metaGRS had higher C-index (C=0.623, 95% CI 0.615–0.630) than any of the individual conventional risk factors, with the second-best factor being hypertension at baseline (C=0.590, 95% CI 0.583–0.598). A model combining the four conventional risk factors had only slightly better performance (C=0.639, 95% CI 0.632–0.647) than the metaGRS individually. Combining the metaGRS with all four conventional risk factors led to a model with C-index of 0.676 (95% CI 0.669–0.684), an increase of 3.7% over the model consisting of the four conventional risk factors.

To investigate the potential role of the metaGRS in earlier life genetic screening, we compared the sex-stratified cumulative incidence of CAD across quintiles of the metaGRS (Figure 3). In UKB men, we observed that CAD risk in the highest metaGRS quintile began exponentially increasing shortly after age 40, reaching a threshold of 10% cumulative risk by 61 years of age (Figure 3). By comparison, CAD risk for men in the lowest metaGRS quintile did not begin increasing until age 50 and on average did not reach 10% by the censoring age of 75. In UKB women, the metaGRS results were similar but delayed given the lower absolute CAD risk overall compared to men. For women in the highest metaGRS quintile, CAD risk began increasing at age 49 and reached 10% at age 75; while women in the lowest metaGRS quintile were at extremely low levels of risk, reaching 2.5% CAD risk by the censoring age of 75. There was no evidence for a statistical interaction of the metaGRS with sex. Overall, on average UKB individuals in the top metaGRS quintile were at 4.17-fold (95% CI 3.97–4.38) higher hazard of CAD than those in the bottom metaGRS quintile (Figure 3).

**Figure 3:**
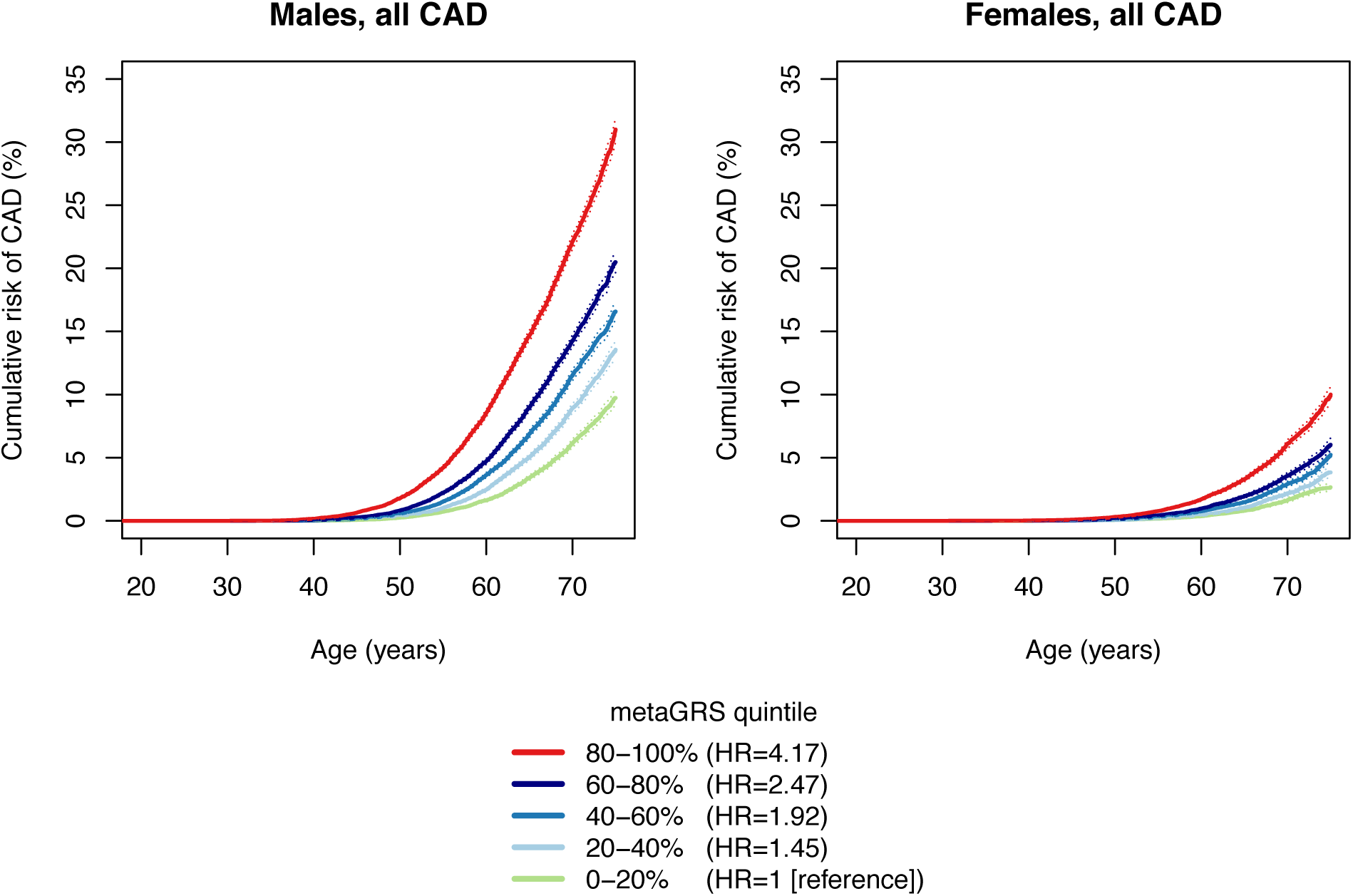
Cumulative incidence of CAD by quintiles of metaGRS in men and women

We next assessed the differences in incident CAD risk across metaGRS quintiles when combined with conventional risk factors (current smoking, diabetes, high blood pressure, and high BMI) individually (**Supplementary Figures 4–7**) or as an unweighted score, the number (0–4) of conventional risk factors per individual (Figure 4). Broadly, the patterns were similar across all the analyses. Genomic risk and lifestyle/clinical factors combined to increase risk in both men and women; however, in most instances this was additive rather than interactive. In Cox regression models of incident CAD, adjusting for current smoking, diagnosed diabetes, hypertension, log BMI, genotyping array, and 10 genetic PCs, there was no strong evidence of statistical interactions between the metaGRS and either diabetes (P=0.051 for interaction), smoking (P=0.086 for interaction), or hypertension (P=0.85 for interaction), but there was some evidence for interaction with log BMI (HR=0.85, 95% CI 0.76–0.5, P=0.0037). From a clinical perspective, it was notable that men in the highest metaGRS quintile who had no conventional risk factors still reached 10% cumulative incidence of CAD by age 69, with a similar cumulative incidence as men in the lowest metaGRS quintile who had 2 or more conventional risk factors (Figure 4). Men in the highest metaGRS quintile and with 3 or more conventional risk factors were at extremely high levels of CAD risk, reaching the 10% threshold by age 48. Approximately 82% of women did not reach 10% CAD risk before age 75, even if they had 2 conventional risk factors, due to compensation by low or moderate metaGRS risk. Even amongst women in the highest metaGRS quintile, only those with 2 or more conventional risk factors achieved 10% risk before age 75 (Figure 4).

**Figure 4:**
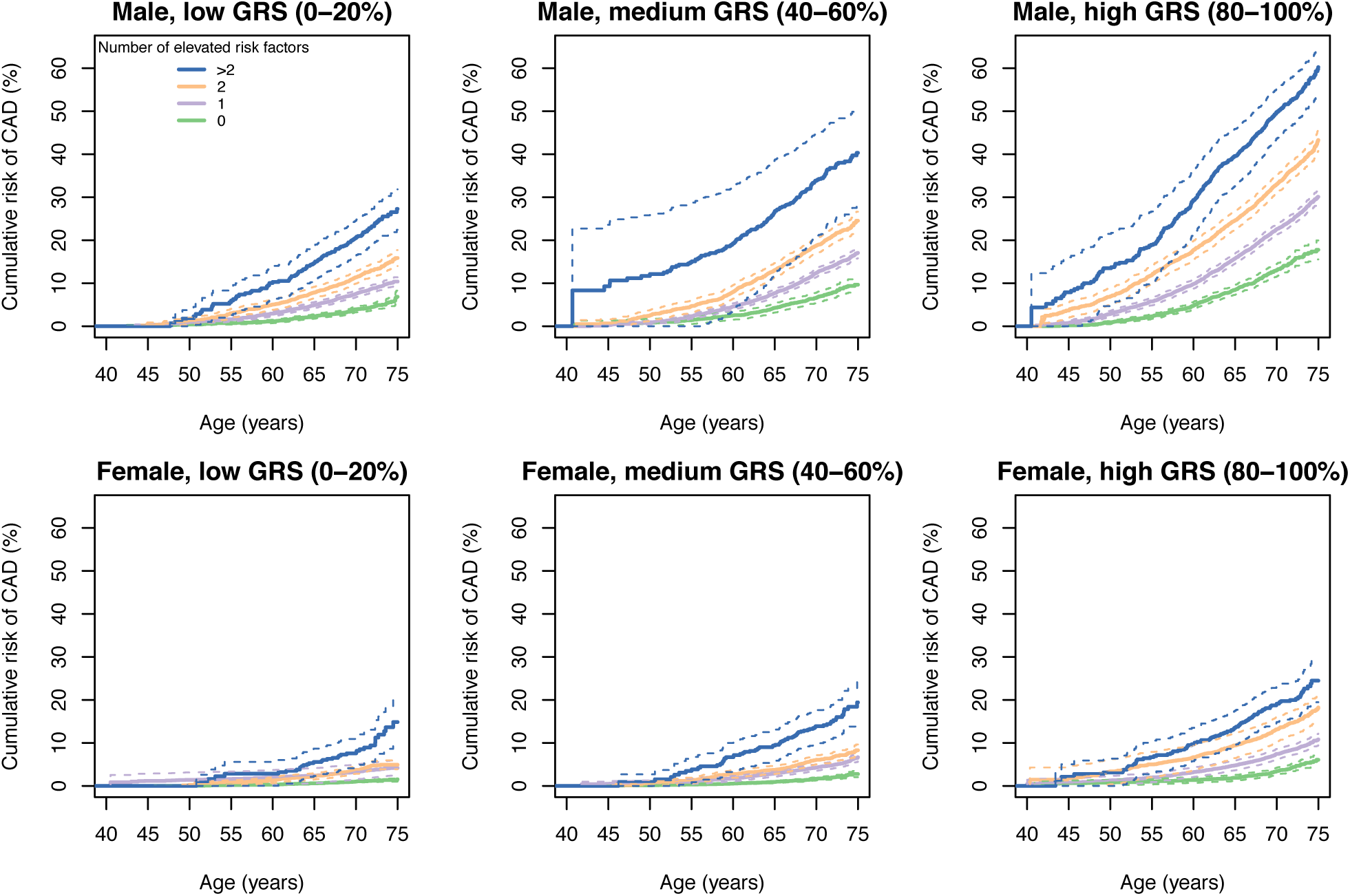
Cumulative incidence of CAD for increasing numbers of conventional risk factors stratified by metaGRS quintile

To assess the impact of use of treatments (lipid lowering and anti-hypertensive medication) that have been proven to lower CAD risk on the performance of the metaGRS, we analysed the predictive capacity of the metaGRS for incident CAD in those taking one or both of these classes of drugs at baseline. The hazards ratios for each s.d. in GRS were reduced but not negated by these therapies, with HRs of 1.44 (95% CI 1.40–1.48), 1.46 (95% CI 1.42–1.50) and 1.42 (95% CI 1.37–1.47) for those individuals on lipid lowering, anti-hypertensives treatments or both treatments, respectively. Accordingly, the HRs between those in the top versus bottom metaGRS quintiles were also reduced but remained substantial with HRs of 2.71 (95% CI 2.47–2.98), 2.81 (95% CI 2.56–3.09), and 2.55 (95% CI 2.28–2.86), for those individuals on lipid lowering, anti-hypertensives treatments or both treatments, respectively (Figure 5).

**Figure 5:**
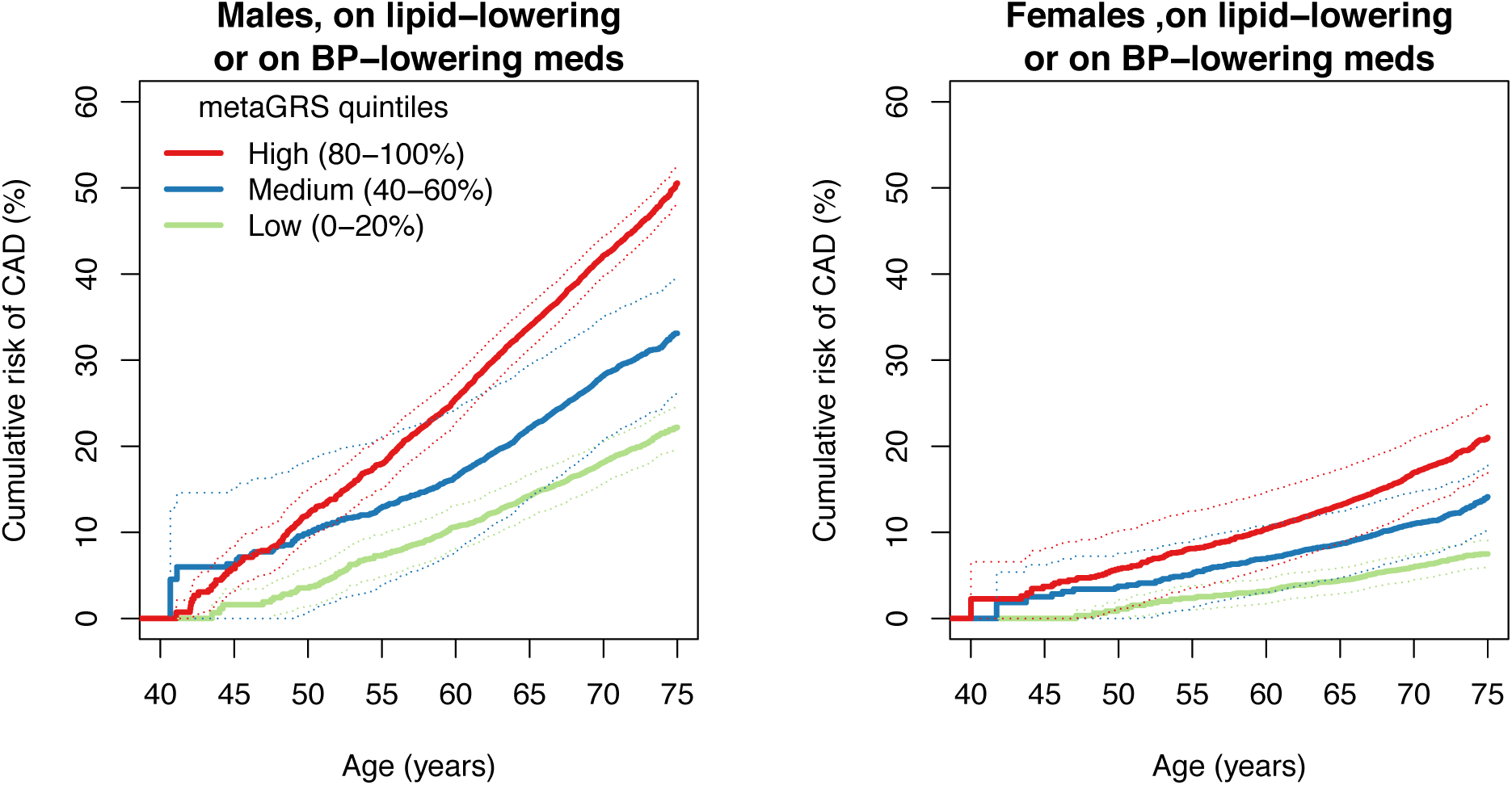
Cumulative incidence of incident CAD within individuals on lipid-lowering or BP-lowering medication at assessment.

## Discussion

Using data from almost half a million people across the UK, we demonstrate that a combined genomic risk score (metaGRS) built from summary statistics of the largest previous genome-wide association studies of CAD performs better than any other individual GRS based on selected SNPs and provides substantial stratification for individual risk of developing CAD. The metaGRS is largely independent of established risk factors for CAD and improves risk prediction using established factors alone or in combination. Importantly, the metaGRS can identify both individuals who are at high risk of premature CAD as well as those who are unlikely to ever reach a life-long risk level requiring intervention. The findings indicate that treatment of modifiable risk factors in those at high risk can partially offset genomic risk, while also highlighting the need to develop additional approaches to address the residual risk. The metaGRS provides valuable additional predictive information at any age, however its unique property of being able to distinguish markedly different lifelong trajectories of individual risk at an early age, before atherosclerosis is initiated, provides the possibility of true primary prevention in those at increased genomic risk, and a potential paradigm shift in how we evaluate risk of and prevent CAD.

Our construction of the metaGRS leveraged the strengths of previous genetic association studies to provide greater predictive power and generalisability than any previous genetic risk score. The metaGRS was stronger than any conventional risk factor available for CAD, largely independent of these risk factors, and substantially increased the predictive power of models combining conventional risk factors. If substantiated once a full set of conventional risk factors has been examined, the metaGRS will have major ramifications for CAD screening, including both identification of individuals at high CAD risk who may benefit from earlier intervention(s) and/or more intensive screening with traditional clinical risk factors, and identification of those individuals at exceptionally low CAD risk who may not reach a clinically relevant level of risk before age 75. Our findings suggest that men in the highest metaGRS quintile, regardless of the number of traditional clinical risk factors, would likely benefit from more intensive interventions; indeed, those with one or more conventional risk factors would likely benefit from statin prescription at an early age. Similar suggestions have been made for Caucasian individuals at high polygenic risk where, using a GRS optimised on a large subset of the UKB, they appeared to have levels of coronary disease risk on par with the risk conferred by familial hypercholesterolemia ^13^. For risk stratification it was notable that approximately 80% of women (i.e. those not in the top metaGRS quintile) could effectively receive minimal screening for traditional risk factors before age 75. Under this scenario, health systems may benefit from more efficient deployment of resources away from individuals at low genomic CAD risk.

The finding that increased genetic risk can at least partly be attenuated by lipid lowering and/or anti-hypertensive medication suggests a potential immediate clinical value to identifying individuals at high metaGRS risk. Those individuals at high genetic risk may gain maximally from early initiation of these therapies and provide more cost-effective primary prevention ^16^. However, the finding that even for individuals on these medications at baseline, the metaGRS can still stratify those at increased risk of CAD, emphasises the need to develop further therapies to realise the full potential of early genetic risk stratification.

The clinical implementation of the metaGRS is straightforward. Each individual’s DNA can be run on one of many common genome-wide genotyping arrays which, together with quality control and genotype imputation, can be combined with a list of genetic variants and their corresponding weights; a simple algorithm then calculates a metaGRS score for that individual. When compared to a large reference group from a similar population, such as the UK Biobank, the individual’s genomic risk of CAD can be determined. Importantly, the genotyping has a one-time cost (approximately US$50 at current prices) and can be used to calculate updated genetic risk scores for CAD as further more powerful association data emerges or, indeed, risk scores for other diseases. To facilitate future development and translation, we have made the metaGRS freely available ^24^.

There are several limitations to our study. Importantly, the UKB does not yet have lipid and other biochemical data available, thus the relationship between the metaGRS and lipids or traditional clinical risk scores (e.g. Framingham Risk Score, QRISK, etc) could not be assessed. However, previous studies have investigated the correlation and added value of genetic risk to clinical risk scores, finding significant improvements in C-index, hazard ratios, and reclassification indices ^3,8^. Another limitation is that the cumulative genomic risk of CAD is likely an underestimate of the population-level lifetime risk due to the likelihood that more individuals at higher genetic risk are more likely to have died from CAD prior to enrolment. Similarly, to avoid reverse causation, incident CAD analyses necessitated the exclusion of individuals with prevalent CAD; however, this also preferentially removed those with early CAD onset and high genomic risk. On the other hand it should be noted that UKB participants are healthier than the general UK population ^25,26^ which could affect the generalisability of the findings. Reverse causation concerns also limited our ability to assess the effect of medication versus non-medication in individuals at high metaGRS risk, since, without blind randomisation, those on medication are already at higher CAD risk. While the UKB captures much of the ethnic diversity of Western Europe, the proportion of non-Caucasian individuals is small (< 5%). We did not exclude these individuals from our analysis, however larger sample sizes and other cohorts will be necessary to undertake meaningful ethnicity-specific analyses. Therefore, the performance and utility of metaGRS in other ethnic populations remains to be determined. Finally, the UKB currently has limited follow-up (median of 6.2 years), therefore both the assessment of clinical risk scores and public health modelling of the metaGRS are important areas for future studies.

In conclusion, our results show that an individual’s genomic risk of CAD, which together with sex is set before birth in germline DNA, is a stable risk factor which can provide the most advanced warning of disease. While genetics is not destiny for CAD, advances in genomic prediction have brought the long history of CAD risk screening to a critical juncture, where we may now be able to predict, plan for, and possibly avoid a disease with substantial morbidity and mortality.

## Acknowledgements

We are grateful to UK Biobank for access to data to undertake our study. This study was supported by funding from National Health and Medical Research Council (NHMRC) grant APP1062227. Supported in part by the Victorian Government’s OIS Program. M.I. was supported by an NHMRC and Australian Heart Foundation Career Development Fellowship (no. 1061435). G.A. was supported by an NHMRC Early Career Fellowship (no. 1090462). N.J.S., C.P.N. and B.K. are supported by the British Heart Foundation and N.J.S. is a NIHR Senior Investigator. R.S.P. is supported by the British Heart Foundation (FS/14/76/30933). The MRC/BHF Cardiovascular Epidemiology Unit is supported by the UK Medical Research Council [MR/L003120/1], British Heart Foundation [RG/13/13/30194], and UK National Institute for Health Research Cambridge Biomedical Research Centre. J.D. is a British Heart Foundation Professor and NIHR Senior Investigator.

